# Myogenic dysregulation underlies tongue overgrowth in Beckwith-Wiedemann syndrome

**DOI:** 10.64898/2026.05.18.725925

**Authors:** Elisia D. Tichy, Anna T. Nguyen, Mariah A. Byrne, Rose D. Pradieu, Gavriela Kalish-Schur, Mara Fallon, Snehal Nirgude, Darryl Kinnear, Harry P. Kozakewich, Jennifer M. Kalish

## Abstract

Macroglossia is a clinically significant feature of Beckwith-Wiedemann syndrome (BWS), but the cellular basis of tongue overgrowth remains poorly defined. Here, using pediatric tongue specimens from molecularly defined BWS subtypes and age-matched nonBWS controls, we show that BWS macroglossia is characterized by skeletal muscle fiber hypertrophy rather than increased fiber number. This phenotype is not explained by expansion or increased proliferation of satellite cells *in situ*, and prospectively isolated tongue satellite cells do not exhibit enhanced proliferation under growth conditions *in vitro*. Instead, BWS progenitors adopt distinct differentiation-associated regulatory states. IC2 loss of methylation cells sustain proliferative activity during differentiation and form enlarged myotubes, consistent with a cell-autonomous hypertrophic program. In contrast, pUPD11 cells display activation of NOTCH signaling and progenitor-associated programs, together with attenuated progression toward terminal myogenic differentiation. These findings identify skeletal muscle hypertrophy as a core tissue-level feature of BWS macroglossia and reveal that epigenetically defined BWS subtypes engage divergent myogenic programs that converge on a shared hypertrophic tissue phenotype. Together, these data define subtype-specific myogenic states in a rare human disease tissue and provide a framework for understanding how distinct epigenetic changes can produce a common overgrowth phenotype.

**Highlights:**

- BWS macroglossia is associated with skeletal muscle fiber hypertrophy, not fiber hyperplasia
- Tongue satellite cell abundance and proliferation are not increased *in situ* in BWS
- IC2 loss of methylation cells sustain proliferation during differentiation and form enlarged myotubes
- pUPD11 cells show enhanced NOTCH signaling and a constrained myogenic state

**In brief:** Tichy *et al.* show that Beckwith-Wiedemann syndrome macroglossia is driven by skeletal muscle hypertrophy and that distinct BWS molecular subtypes engage different myogenic regulatory programs. IC2 loss of methylation cells sustain proliferation during differentiation, whereas pUPD11 cells exhibit NOTCH-associated restraint of myogenic progression.

## Introduction

Beckwith-Wiedemann syndrome (BWS) is an imprinting disorder characterized by somatic overgrowth, developmental abnormalities, and increased cancer risk ^1^. Most cases arise from epigenetic or genetic alterations affecting chromosome 11p15, a locus containing two imprinting control regions (IC1 and IC2) that regulate dosage-sensitive growth pathways and include key growth regulatory genes such as *IGF2* and *CDKN1C* ^2^. The most common molecular subtypes are IC2 loss of methylation (IC2 LOM) and paternal uniparental disomy of chromosome 11 (pUPD11) ^3^. Notably, pUPD11 produces combined dysregulation of both imprinting domains, resulting in IC1 gain of methylation together with IC2 loss of methylation within affected cells. Less frequently, BWS arises from isolated IC1 gain of methylation, pathogenic variants in *CDKN1C*, or structural abnormalities of the 11p15 region ^3^. Although these molecular subtypes share disruption of growth-regulatory pathways, they are associated with distinct clinical patterns and tissue-specific overgrowth phenotypes.

Macroglossia, or pathological tongue enlargement, is among the most recognizable and clinically significant manifestations of BWS ^4^. Excessive tongue overgrowth can lead to airway obstruction, feeding difficulties, and speech impairment, often requiring surgical tongue reduction during early childhood ^5^. Clinical tools such as the Beckwith-Wiedemann syndrome Index of Macroglossia (BIG) score have improved risk stratification for surgical intervention ^6^. However, the biological mechanisms underlying tongue overgrowth remain poorly understood. Importantly, patients requiring surgical tongue reduction are disproportionately enriched for the IC2 LOM and pUPD11 subtypes, suggesting that dysregulation of these imprinting domains plays a central role in the development of macroglossia.

The tongue is a complex organ composed primarily of skeletal muscle, together with connective tissue, vasculature, nerves, and specialized epithelial structures. During embryogenesis, intrinsic tongue musculature arises from migratory myogenic progenitors derived from occipital somites ^7^. These progenitors express the paired box transcription factors PAX3 and PAX7, which are essential regulators of skeletal muscle lineage specification and development ^8,9^. In postnatal tissue, PAX3- and PAX7-expressing satellite cells function as muscle stem cells that support skeletal muscle growth and regeneration through coordinated cycles of proliferation, differentiation, and fusion into multinucleated muscle fibers ^10,11^. Dysregulation of these processes is a well-established driver of skeletal muscle hypertrophy and disease ^12^.

Previous case reports have suggested conflicting explanations for BWS macroglossia, proposing either skeletal muscle hypertrophy (enlarged fibers) or hyperplasia (increased fiber number) ^13^. However, these studies were limited by small sample sizes, adult tissue analyses, or the absence of appropriate pediatric controls. Moreover, little is known about how the distinct molecular subtypes of BWS influence the cellular mechanisms underlying tongue overgrowth. Because IC2 LOM and pUPD11 share disruption of the IC2 imprinting domain but differ in their broader epigenetic context, these subtypes provide a unique opportunity to examine how related imprinting defects may produce distinct cellular outcomes within the same tissue.

Here, we analyze pediatric tongue tissue from molecularly defined BWS patients undergoing tongue reduction surgery and compare these samples with age-matched nonBWS pediatric controls from autopsy. We show that macroglossia in both IC2 LOM and pUPD11 is associated with skeletal muscle fiber hypertrophy rather than increased fiber number. Despite this shared tissue phenotype, satellite cell abundance and proliferative activity are not increased *in situ*. Functional analyses of isolated satellite cells instead reveal subtype-specific differentiation behaviors, with IC2 LOM cells exhibiting sustained proliferation during differentiation and enlarged myotube formation, whereas pUPD11 cells display enrichment of NOTCH signaling and progenitor-associated programs. These findings demonstrate that distinct epigenetically defined myogenic programs underlie tongue overgrowth in BWS and provide a mechanistic framework for understanding subtype-specific myogenic dysregulation.

## Results

### BWS-associated macroglossia is characterized by skeletal muscle fiber hypertrophy

We previously reported that tongues from patients with IC2 LOM are characteristically wider, whereas pUPD11 tongues are narrower and thicker, based on morphometric analyses ^4^ (Figure 1A). To define the cellular basis of macroglossia in the two most prevalent BWS subtypes, we compared tongue specimens from IC2 LOM and pUPD11 reduction surgeries with nonBWS pediatric autopsy controls. Sample characteristics are presented in Figure S1. Histological analyses revealed pronounced enlargement of skeletal muscle fibers in both BWS subtypes relative to controls (Figure 1B). Quantification of fiber cross-sectional area demonstrated that control tongues contained smaller muscle fibers than either IC2 LOM or pUPD11 tongues (Figure 1C and Figure S2). Fiber size distribution analysis similarly showed that control tongues contained smaller fibers than either IC2 LOM or pUPD11 tongues (Figure 1D and 1E). In contrast, fiber size distributions were comparable between IC2 LOM and pUPD11 samples (Figure 1F). Together, these findings identify skeletal muscle fiber hypertrophy as the principal tissue-level feature of BWS-associated macroglossia.

**Figure 1.**
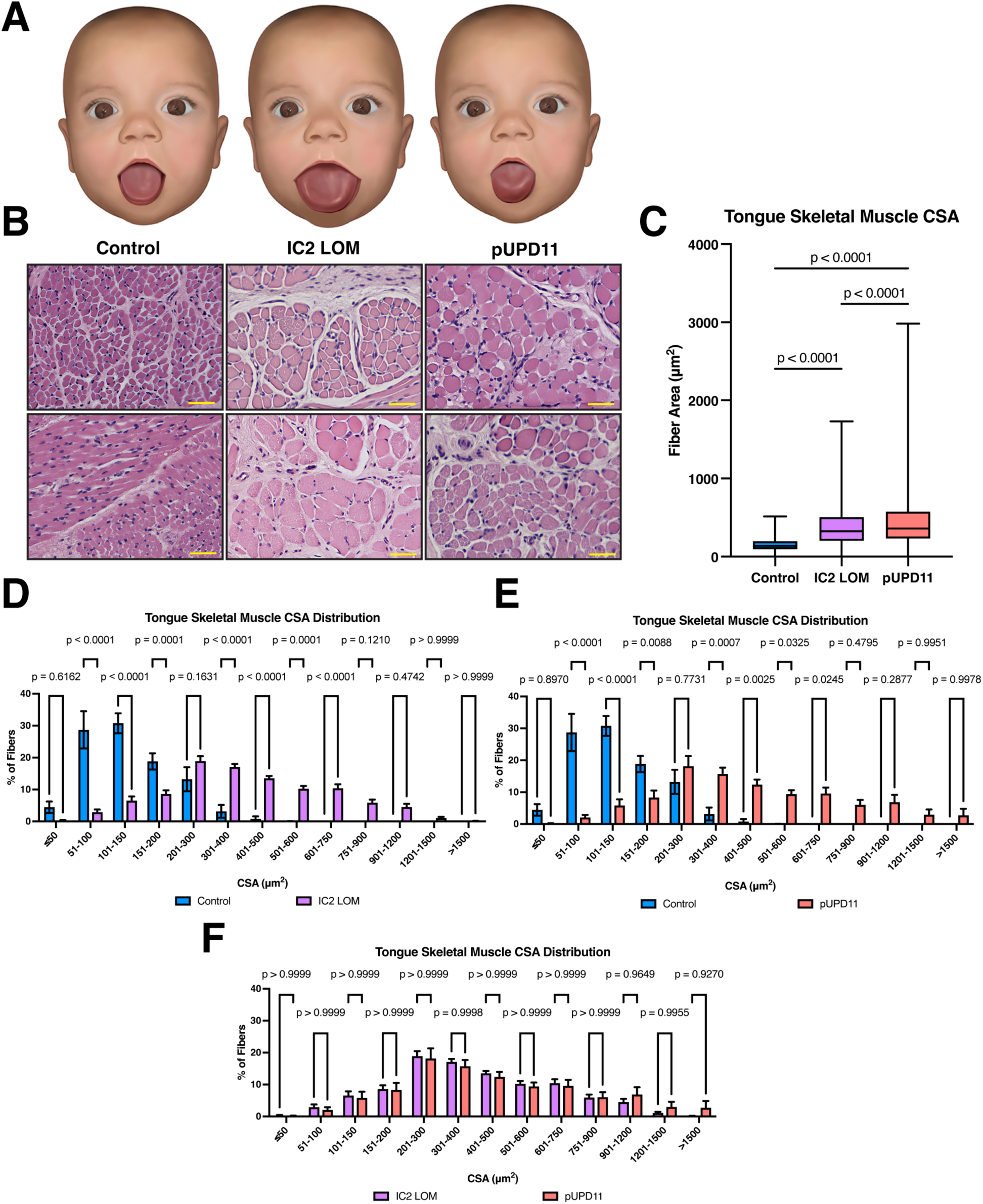
BWS macroglossia is associated with skeletal muscle fiber hypertrophy. (A) Schematic renderings of control (nonBWS), IC2 loss of methylation (IC2 LOM), and paternal uniparental disomy of chromosome 11 (pUPD11) tongues based on clinical observations. (B) Representative hematoxylin and eosin–stained sections from control, IC2 LOM, and pUPD11 tongues. Scale bars: 50 μm. (C) Box-and-whisker plots showing muscle fiber cross-sectional area (CSA) measured from 6 control, 25 IC2 LOM, and 10 pUPD11 patients. At least 1,000 muscle fibers were analyzed per patient. Data were analyzed by one-way ANOVA; P values are indicated. (D–F) Frequency distributions of muscle fiber CSA comparing control versus IC2 LOM (D), control versus pUPD11 (E), and IC2 LOM versus pUPD11 (F) tongue samples. Data were analyzed by two-way ANOVA; P values are shown.

### Satellite cell abundance and proliferative activity are not increased in BWS tongues

Skeletal muscle growth is regulated by satellite cells, which function as resident muscle stem cells. During development and postnatal growth, these cells undergo regulated cycles of proliferation and differentiation to generate skeletal muscle fibers ^14^. Because hypertrophy of skeletal muscle fibers was observed in BWS tongues, we examined whether macroglossia was associated with changes in satellite cell abundance or proliferative activity. Immunostaining of tongue sections revealed similar numbers of PAX3+ and PAX7+ cells across control, IC2 LOM, and pUPD11 samples (Figure S3A, S3B, S3D, and S3E). Co-staining with the proliferation marker KI67 ^15^ also showed no significant differences in satellite cell proliferation between groups (Figure S3C and S3F). Expression of MYOD, a marker of myogenic activation and myoblast identity ^16^, was variable across samples but did not differ significantly between experimental groups (Figure S3G and S3H). Enumeration of KI67 positivity within MYOD+ cells likewise revealed no significant differences (Figure S3I). These findings suggest that satellite cell abundance and proliferative activity are not increased in BWS tongues at the time of surgical resection.

### Satellite cells from BWS tongues do not exhibit increased proliferation in growth-permissive conditions *in vitro*

To determine whether intrinsic proliferative differences might emerge under controlled conditions, we isolated live satellite cells from cryobanked tongue tissue using established human satellite cell markers CD56 and CD82 ^17,18^. Inflammatory and epithelial populations were excluded by gating out CD45+ and CD324+ cells, resulting in a clearly defined satellite cell population (Figure S4A). Purified cells were cultured under growth-permissive conditions for four days and assayed for proliferation using EdU incorporation (Figure S4B). Across all groups, satellite cells displayed comparable proliferative capacity, with no statistically significant differences between control, IC2 LOM, and pUPD11 samples (Figure S4C). Although BWS-derived cells showed modest increases in proliferative potential, these differences were insufficient to account for the pronounced skeletal muscle hypertrophy observed *in vivo*.

### IC2 LOM satellite cells form enlarged myotubes during differentiation

To determine whether altered differentiation or fusion contributes to muscle hypertrophy, satellite cells were induced to differentiate *in vitro* (Figure 2A). Robust myotube formation was observed across all groups by Day 10 after plating (Figure S5A). Expression of the early differentiation marker MYOGENIN did not differ significantly between control and BWS samples (Figure S5B and S5C), indicating that initiation of the differentiation program was not altered. However, staining for myosin heavy chain (MF20) revealed marked morphological differences between experimental groups. IC2 LOM cultures exhibited enlarged myotubes with clustered nuclei, a phenotype that was less prominent in pUPD11 cultures and absent in controls (Figure 2B). Fusion indices were not significantly different between groups (Figure 2C), but quantitative analysis demonstrated a significant increase in myosin heavy chain-positive area in IC2 LOM cultures compared with controls (Figure 2D). These findings indicate that IC2 LOM satellite cells display altered differentiation behavior that recapitulates the hypertrophic phenotype observed *in vivo*.

**Figure 2.**
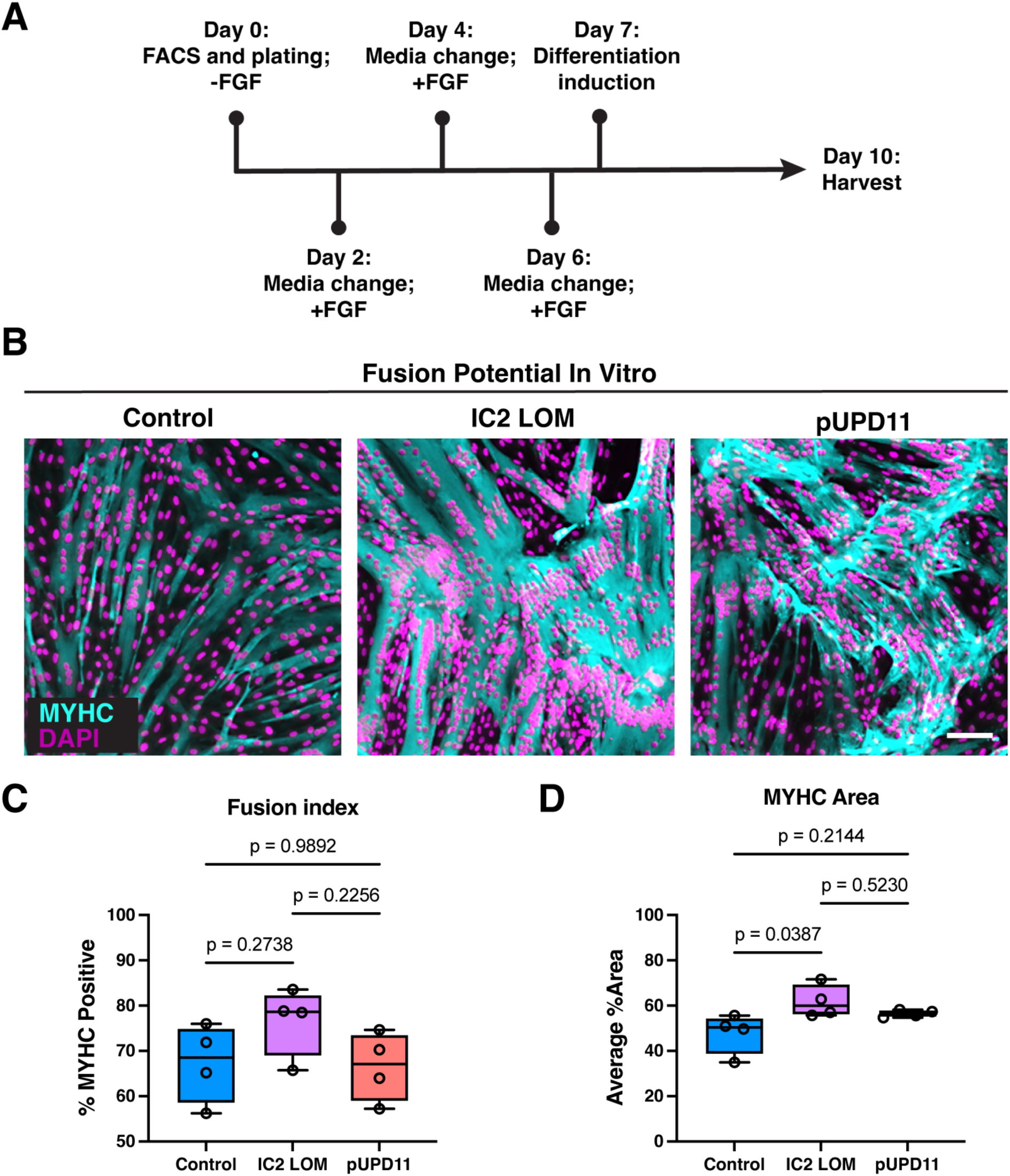
IC2 LOM tongue satellite cells form enlarged myotubes during differentiation. (A) Experimental timeline for satellite cell plating and assay mechanics. (B) Representative images of FACS-isolated myogenic cells from control, IC2 LOM, and pUPD11 samples differentiated for 3 days and immunostained for myosin heavy chain (MyHC; MYH4; MF20; cyan) to visualize myotube formation. Scale bar: 100 μm. (C) Quantification of fusion index for control, IC2 LOM, and pUPD11 cells. Fusion index was calculated as the number of nuclei within MyHC⁺ myotubes containing two or more nuclei divided by the total number of nuclei. (D) Quantification of total MyHC⁺ area per field in control, IC2 LOM, and pUPD11 cultures. Data are shown as box-and-whisker plots and were analyzed by one-way ANOVA. Each data point represents a sample derived from a unique patient.

### RNA sequencing identifies subtype-specific signaling programs during myogenic differentiation

To identify molecular pathways underlying the observed subtype-specific differentiation behaviors, we performed very low-input bulk RNA sequencing on satellite cells collected at Day 9 post-plating (Day 2 of differentiation). This time point was selected to capture transcriptional differences preceding the overt morphological divergence observed at Day 10. Principal component analysis (PCA) revealed clear separation between control and BWS samples, whereas IC2 LOM and pUPD11 samples partially overlapped (Figure 3A). Pathway enrichment analysis comparing IC2 LOM cells with controls identified proliferative signaling pathways, including CDK4 and CDK6 activity, as the most significantly enriched pathways (Figure 3B). Additional enriched pathways in IC2 LOM were predominantly associated with collagen organization and extracellular matrix (ECM) remodeling. In contrast, pUPD11 cells exhibited enrichment of pathways related to NOTCH signaling, muscle fiber formation, and insulin-like growth factor (IGF) transport (Figure 3C). Reciprocal comparisons further highlighted subtype-specific signaling differences. Relative to IC2 LOM cells, control samples were enriched for RUNX- and WNT-associated signaling pathways, as well as G protein-coupled receptor signaling (Figure 3D). Comparisons between control and pUPD11 cells revealed downregulation of pathways related to muscle contraction, ECM organization, and progenitor-associated interactions in pUPD11 cells (Figure 3E). STRING-based network analyses were consistent with these findings. IC2 LOM versus control comparisons identified ECM-centered and pro-proliferative interaction modules, whereas pUPD11 versus control comparisons confirmed enrichment of NOTCH-associated and growth factor-related networks, alongside downregulation of muscle structural and metabolic modules (Figures S6 and S7).

**Figure 3.**
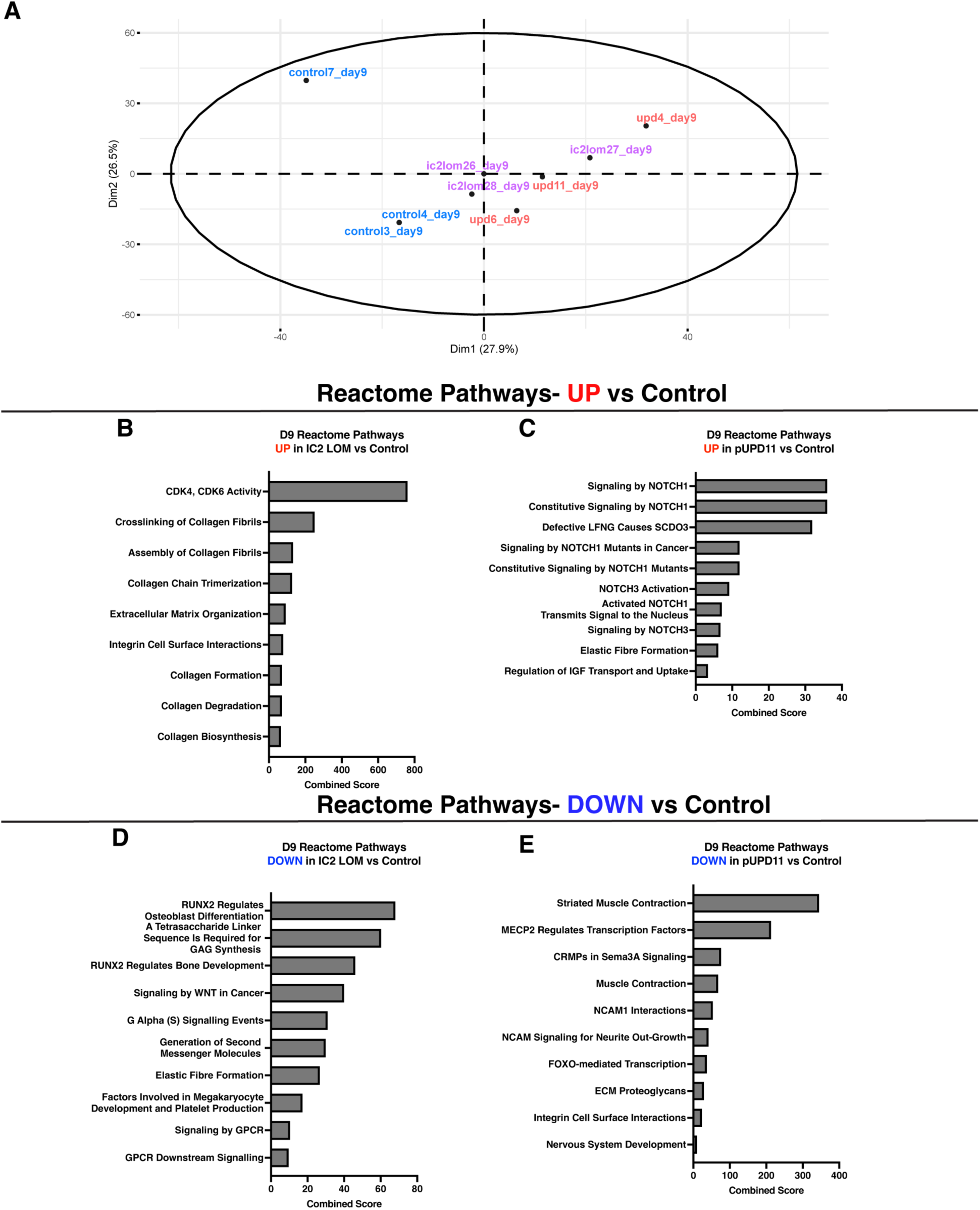
Low-input RNA sequencing reveals subtype-specific transcriptional programs. (A) Principal component analysis (PCA) of satellite cell samples collected at Day 9 post-plating and subjected to RNA sequencing. (B) Reactome pathway enrichment analysis of genes upregulated in IC2 LOM relative to control, performed using Enrichr and ranked by combined score. (C) Reactome pathway enrichment analysis of genes upregulated in pUPD11 relative to control. (D) Pathways enriched in control relative to IC2 LOM samples. (E) Pathways enriched in control relative to pUPD11 samples.

### Divergent NOTCH signaling and myogenic gene expression distinguish BWS subtypes

Given the subtype-specific pathway enrichment identified by RNA sequencing, we next examined key signaling and myogenic gene expression programs to define the regulatory states distinguishing IC2 LOM and pUPD11 cells. To further interrogate the role of NOTCH signaling and myogenic programs, we performed gene set enrichment analysis using curated gene sets associated with NOTCH signaling, skeletal muscle, and myogenesis. In IC2 LOM cells, only a limited number of NOTCH pathway genes were differentially expressed relative to controls, including reduced expression of *HEY1* and *GATA5* and increased *FOXC2* and *SORBS2* (Figure 4A). In contrast, pUPD11 cells exhibited broad activation of NOTCH signaling components, including increased expression of *NOTCH1*, *NOTCH3*, *JAG1*, *DLL1*, *HEY1*, and *TSPAN15* (Figure 4B). Negative regulators of NOTCH signaling, including the E3 ubiquitin ligases *FBXW7* and *NEURL1*, were downregulated (Figure 4B), consistent with sustained pathway activation. Analysis of skeletal muscle differentiation genes further revealed enrichment of progenitor-associated genes including *PAX7* and *SOX8* in pUPD11 cells, whereas control cells were enriched for differentiation-associated genes including *MEF2C* and *KLHL40*, *KLF5*, and *SOX11* ^19–21^ (Figures 4C and 4D). Examination of myogenesis-associated genes further revealed enrichment of IGF-binding proteins, the embryonic muscle marker *TAGLN*, the ECM component *COL15A1*, and the fast-twitch myosin isoform *MYH2* in IC2 LOM cells (Figure 4E). In contrast, pUPD11 cells were enriched for NOTCH1 and AEBP1, a known inhibitor of muscle differentiation ^22^, together with widespread downregulation of sarcomeric genes (Figure 4F). These findings indicate that pUPD11 cells retain a NOTCH-associated progenitor state, whereas IC2 LOM cells progress toward a hypertrophic differentiation program.

**Figure 4.**
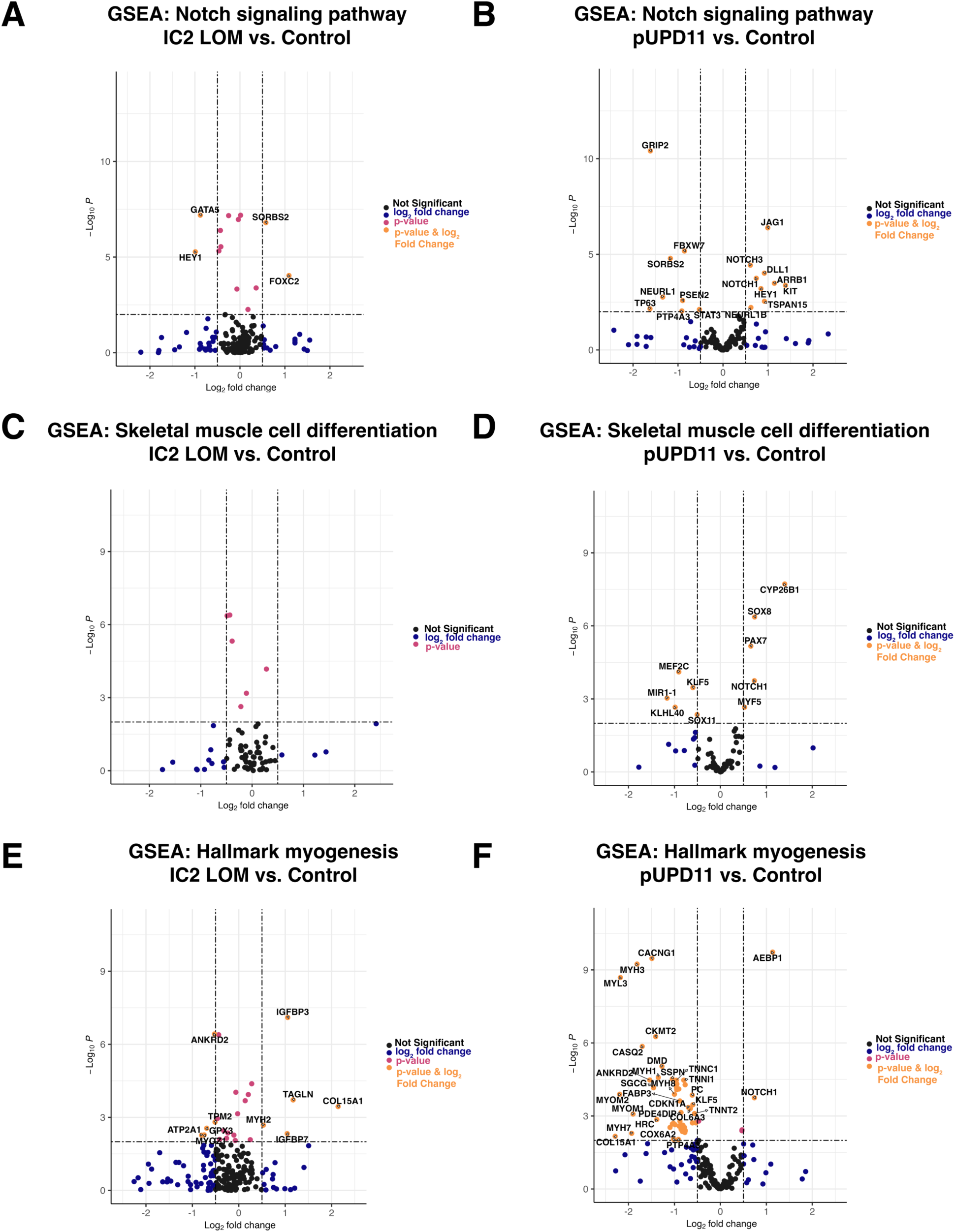
pUPD11 cells retain a NOTCH-associated restraint of myogenic progression. Volcano plots generated from Day 9 RNA sequencing data using curated human gene sets showing differential expression of genes associated with (A) NOTCH signaling in IC2 LOM versus control, (B) NOTCH signaling in pUPD11 versus control, (C) skeletal muscle differentiation in IC2 LOM versus control, (D) skeletal muscle differentiation in pUPD11 versus control, (E) myogenesis in IC2 LOM versus control, and (F) myogenesis in pUPD11 versus control. Black points indicate genes that are neither differentially expressed nor statistically significant. Blue points indicate genes that are differentially expressed but do not meet the significance threshold. Pink points represent genes that are statistically significant but do not meet the predefined threshold for differential expression. Orange points indicate genes that are both differentially expressed and statistically significant.

### IC2 LOM satellite cells sustain proliferation during differentiation

Because RNA sequencing suggested enrichment of proliferative pathways in IC2 LOM cells, we next examined proliferation during myogenic differentiation. Satellite cells were pulsed with EdU at defined intervals during differentiation and co-stained for myosin heavy chain (Figure 5A). At Day 8 post-plating (Day 1 of differentiation), IC2 LOM samples showed a trend toward increased proliferation relative to controls and significantly higher proliferation compared with pUPD11 samples (Figures 5B and 5C). By Day 9, proliferation levels in IC2 LOM and control cultures were similar, whereas pUPD11 cells continued to show reduced proliferative activity (Figures 5D and 5E). By Day 10, proliferation remained elevated in IC2 LOM cultures while control cultures declined to levels similar to those observed in pUPD11 cells (Figures 5F and 5G). Consistent with these findings, IC2 LOM cultures contained a greater number of EdU-positive nuclei within or adjacent to myosin heavy chain-positive myotubes compared with control or pUPD11 cultures (Figures S8A and S8B). These findings support a model in which IC2 LOM satellite cells retain proliferative capacity during late differentiation, thereby increasing nuclear contribution to growing myotubes and promoting hypertrophic muscle formation.

**Figure 5.**
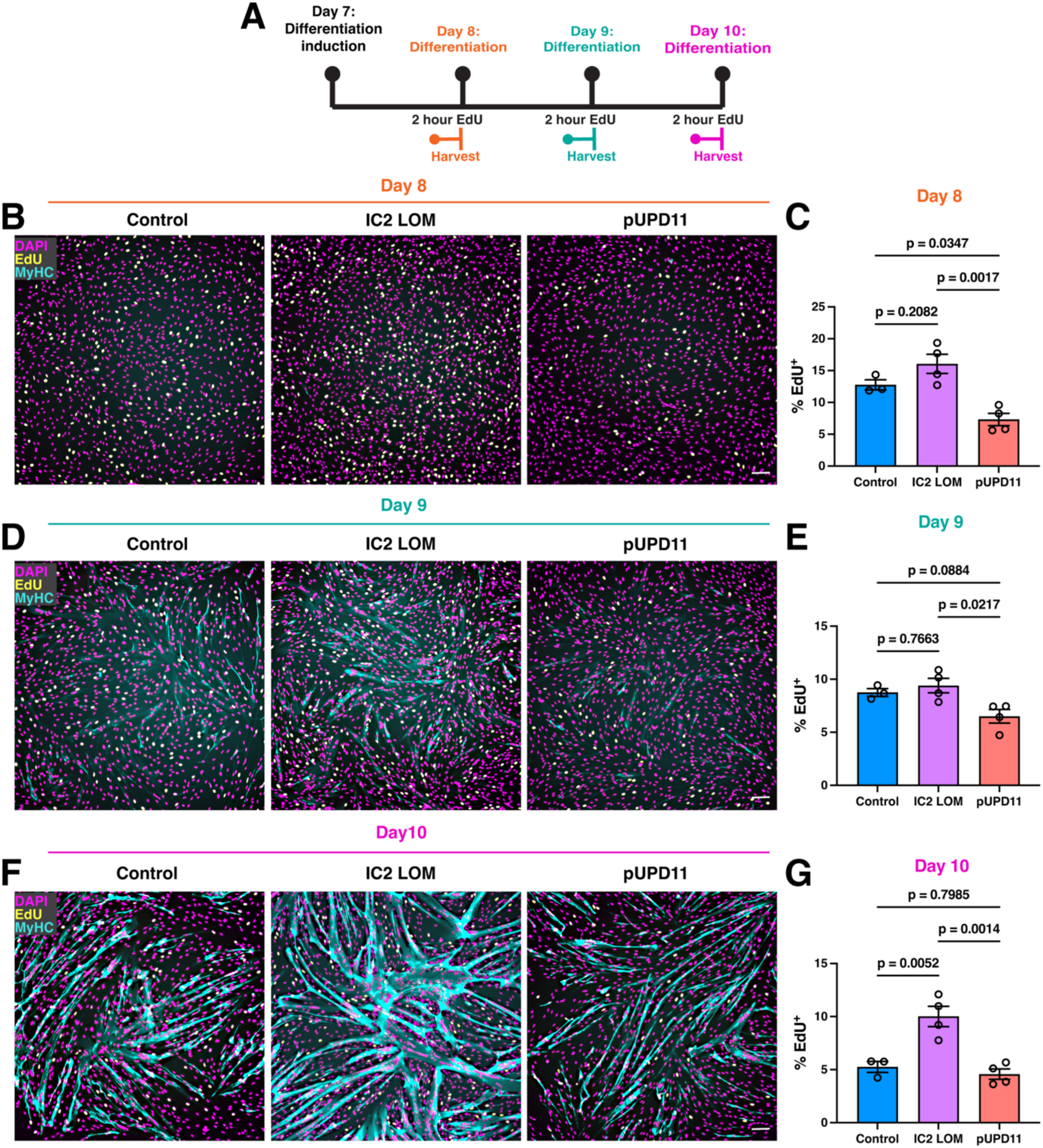
IC2 LOM cells sustain proliferation during differentiation. (A) Schematic of the experimental design for EdU pulse labeling during differentiation. (B, D, F) Representative images of satellite cell cultures from control, IC2 LOM, and pUPD11 samples pulsed with EdU for 2 h and harvested at Day 8 (B), Day 9 (D), or Day 10 (F) of differentiation, followed by co-immunostaining for EdU incorporation and MyHC. Scale bar: 100 μm. (C, E, G) Quantification of the percentage of EdU⁺ nuclei at one (C), two (E), or three (G) days after differentiation induction. Each data point represents cells derived from a unique patient. Data were analyzed by one-way ANOVA; P values are shown.

## Discussion

Macroglossia is one of the most clinically significant manifestations of BWS, yet the biological basis of tongue overgrowth has remained poorly defined. By integrating pediatric tissue analysis, prospective isolation of tongue satellite cells, and low-input transcriptomics, we show that BWS macroglossia is associated with skeletal muscle fiber hypertrophy and that the two major BWS molecular subtypes engage distinct myogenic regulatory programs.

Our histological analyses establish muscle fiber hypertrophy, rather than increased fiber number, as the dominant tissue feature in BWS macroglossia. This clarifies a long-standing uncertainty in the field and provides a more specific framework for understanding how imprinting defects alter tongue growth. Importantly, the hypertrophic phenotype was observed in both IC2 LOM and pUPD11 samples, indicating that different molecular subtypes can converge on a common anatomical outcome.

Despite this shared tissue phenotype, we found no evidence that increased tongue size at the time of surgery is maintained by a larger or more proliferative satellite cell compartment *in situ*. This finding suggests that the initiating overgrowth events likely occur earlier in development, potentially during prenatal or early postnatal myogenesis. The absence of overt *in situ* differences at the time of tissue collection underscores the value of functional *ex vivo* assays to reveal latent subtype-specific cell states.

These assays uncovered a striking mechanistic distinction between the two major BWS subtypes. IC2 LOM satellite cells did not exhibit increased proliferation in growth medium, but during differentiation they formed enlarged myotubes and sustained proliferative activity later than controls. These findings are consistent with a cell-autonomous mechanism for excessive muscle growth and define IC2 LOM cells as adopting a hypertrophic differentiation state. Transcriptomic enrichment of CDK-related pathways provides a plausible mechanistic basis for this phenotype, particularly in the context of reduced CDKN1C dosage associated with IC2 LOM. Enrichment of extracellular matrix-related programs further suggests that altered cell-matrix interactions may reinforce abnormal myogenic behavior.

In contrast, pUPD11 cells were marked by strong NOTCH pathway activation and increased expression of progenitor-associated and quiescence-associated genes such as PAX7 ^23^ and SOX8 ^24^. This state was accompanied by reduced proliferative activity during differentiation and downregulation of genes encoding structural components of differentiated muscle. Because NOTCH signaling is a well-established regulator of satellite cell quiescence and restraint of differentiation ^25–27^, these data support a model in which pUPD11 cells remain in a NOTCH-associated progenitor state and are less intrinsically capable of driving hypertrophic muscle formation under isolated culture conditions.

This distinction is biologically important because IC2 LOM and pUPD11 share IC2 loss of methylation, yet they do not behave equivalently. Our findings suggest that additional dosage changes present in pUPD11, including broader disturbance across 11p15, may shift myogenic cells into a state of persistent NOTCH-associated restraint. As a result, pUPD11-associated macroglossia may depend more heavily on non-cell-autonomous cues from the tissue microenvironment than IC2 LOM-associated disease.

Together, these findings show that distinct epigenetic subtypes of BWS converge on a shared anatomical phenotype through different myogenic mechanisms. IC2 LOM is associated with a cell-autonomous hypertrophic program, whereas pUPD11 is associated with a NOTCH-linked progenitor state that likely requires non-cell-autonomous cues to promote tissue overgrowth in vivo. These results define subtype-specific myogenic dysregulation as a basis for tongue overgrowth in BWS.

## Limitations of the study

Pediatric tongue specimens from molecularly defined BWS patients represent an exceptionally rare resource for studying the cellular mechanisms of disease. Consequently, this study is limited by the rarity of clinically indicated pediatric tongue specimens and pediatric autopsy control specimens, resulting in smaller sample sizes for some functional assays. The transcriptomic analyses were performed using very low-input bulk RNA sequencing and therefore do not resolve cell-state heterogeneity at single-cell resolution. Because tissues were obtained at the time of clinically indicated surgery, our analyses likely capture downstream consequences of earlier developmental perturbations rather than the initiating *in vivo* events. Finally, *in vitro* differentiation assays may not fully recapitulate the extracellular environment of the developing tongue, particularly for pUPD11 samples that may rely on non-myogenic signals.

## STAR Methods

### Key resources table

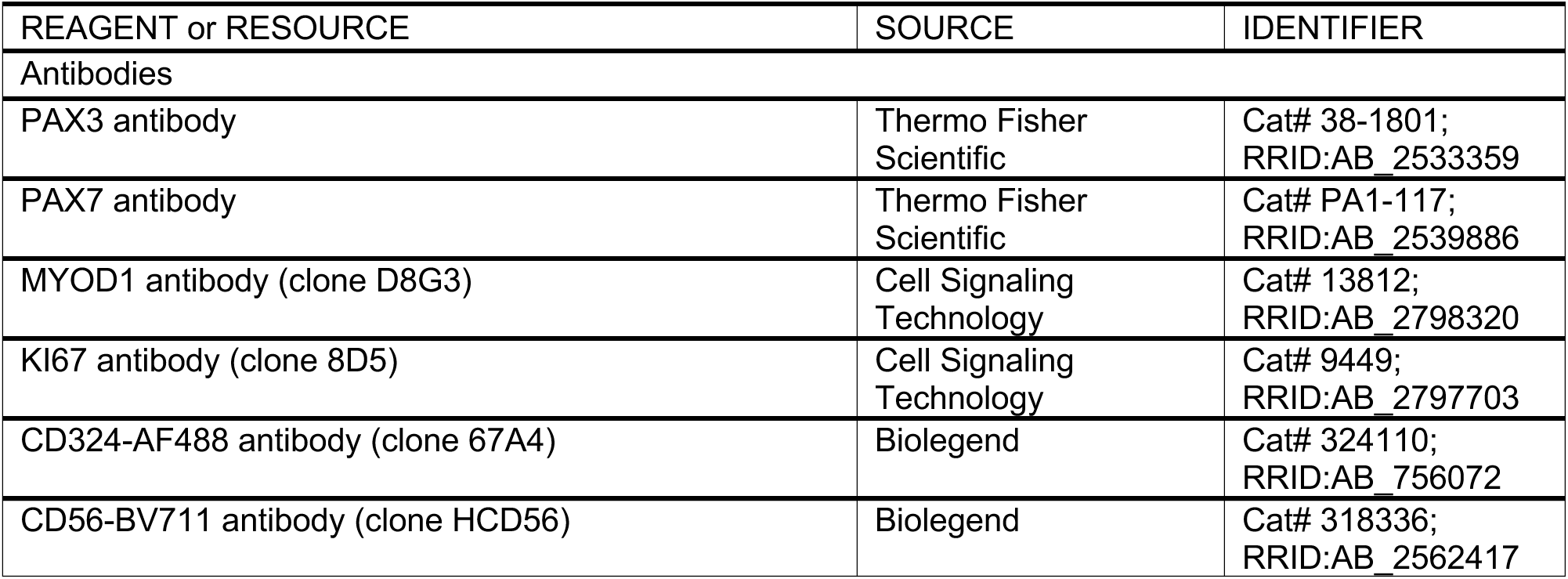
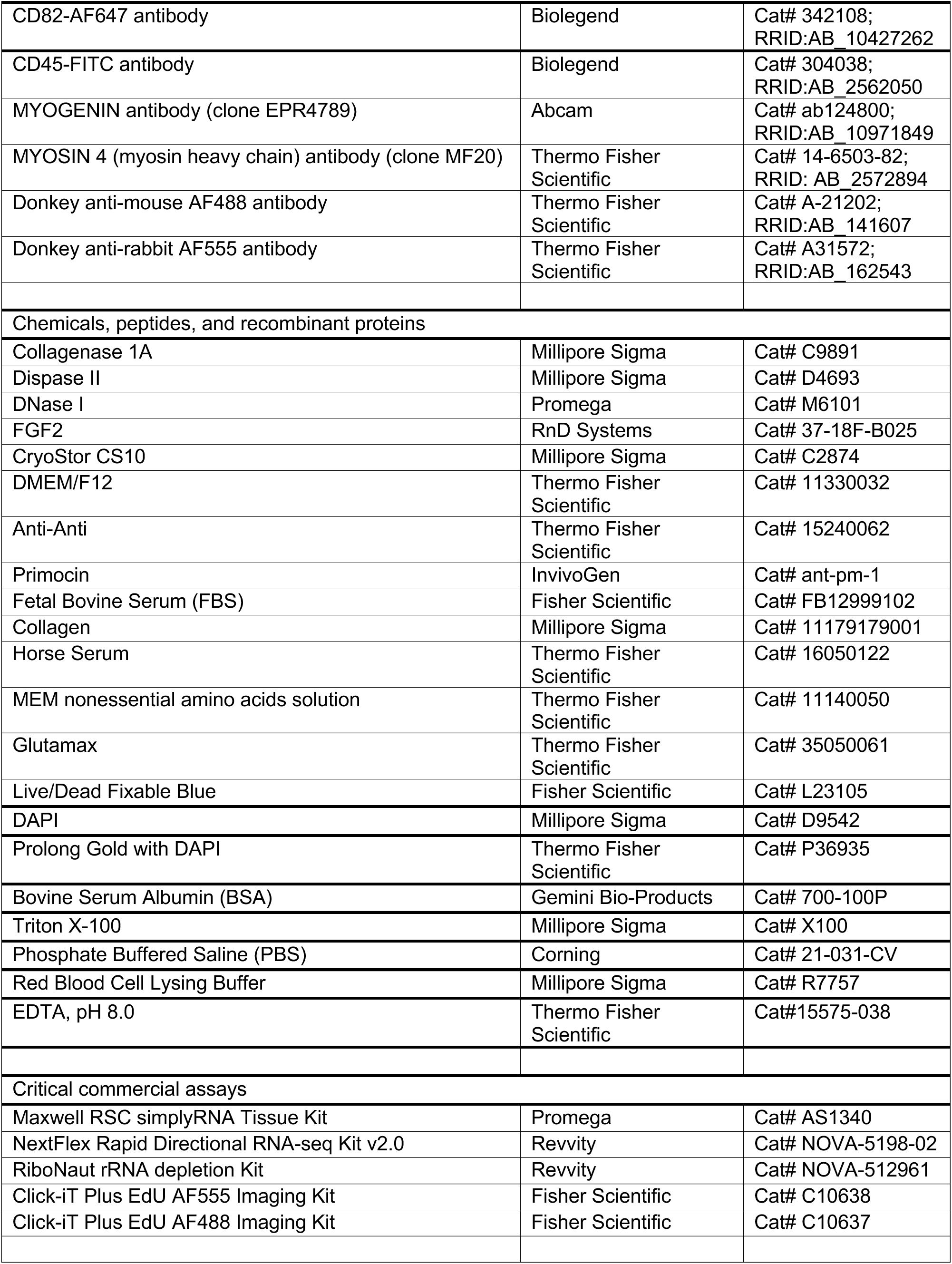
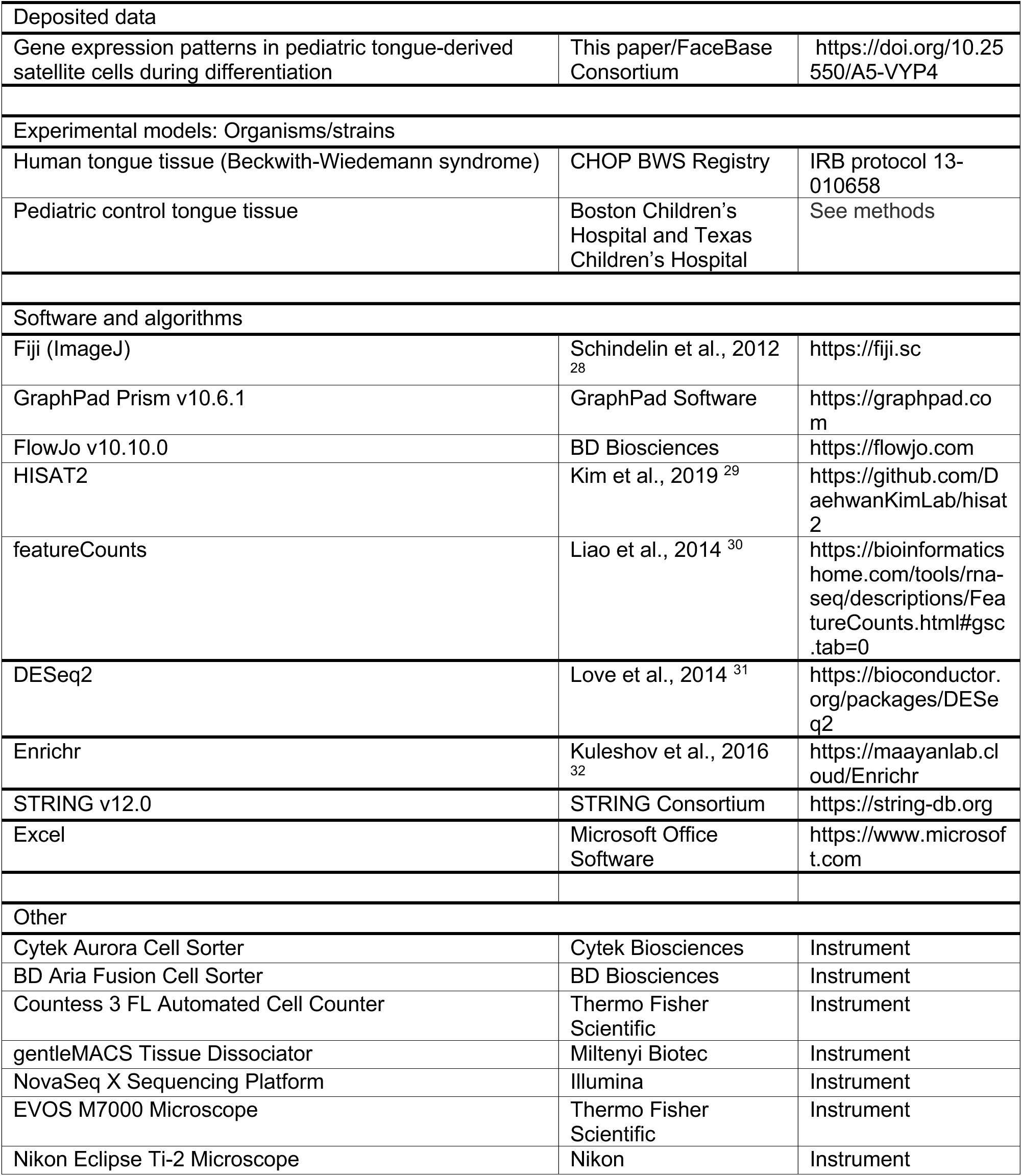

### Resource availability

Further information and requests for resources and reagents should be directed to the lead contact, Jennifer M. Kalish.

### Materials availability

There are restrictions to the availability of patient-derived tongue specimens because of IRB protocol limitations.

### Data and code availability

- Sequencing reads have been deposited in FaceBase and are publicly available as of the date of publication: https://doi.org/10.25550/A5-VYP4
- All numerical data underlying graphs are provided in Supplemental Data File 1.
- This study did not generate custom code.

## Experimental model and study participant details

### Human tissue specimens

All BWS patient samples and associated clinical information were collected through the Children’s Hospital of Philadelphia Beckwith-Wiedemann syndrome registry under institutional review board protocol IRB 13-010658. All patients had molecularly confirmed BWS diagnoses established during clinical evaluation. All experiments were performed in accordance with the Declaration of Helsinki. Written informed consent was obtained for all BWS samples, which were collected at the time of clinically indicated tongue reduction surgery. Samples were selected based on availability, without preference for donor sex.

NonBWS control samples were obtained from pediatric autopsy specimens from children younger than 3 years of age. Control inclusion criteria were absence of macroglossia or microglossia, no history of cancer, and no major facial trauma. Specimens were collected within 24 h of death.

### Tissue processing

Tongue tissue was processed for formalin-fixed paraffin embedding, OCT cryoembedding, and cryobanking. For cryobanking, tissue fragments were washed three times in Hanks’ balanced salt solution containing 2X Primocin, mechanically diced, transferred into CryoStor CS10, slowly frozen in a CoolCell freezing container, and stored in the vapor phase of liquid nitrogen.

## Method details

### Histology and muscle fiber cross-sectional area analysis

Five-micrometer FFPE tongue sections were deparaffinized, rehydrated, and stained with hematoxylin and eosin using standard procedures. Slides were imaged at 10x magnification on an EVOS M7000 microscope. Muscle fiber cross-sectional area was measured in FIJI by manual tracing of individual fibers. For fiber size distribution analyses, bins spanning the full range of fiber sizes were generated in Excel using the data analysis toolpak, and fiber percentages were plotted in GraphPad Prism.

### Immunofluorescence of tongue cryosections

Ten-micrometer cryosections were fixed in 4% paraformaldehyde, permeabilized with 0.5% Triton X-100, and blocked in 3% BSA with 0.1% Triton X-100. Primary antibodies against PAX3, PAX7, MYOD1, and/or KI67 were incubated overnight at 4°C. Appropriate Alexa Fluor-conjugated secondary antibodies were applied for 1 h at room temperature. Wheat germ agglutinin conjugated to Alexa Fluor 647 was included to visualize myofiber boundaries. Coverslips were mounted with ProLong Gold containing DAPI. Images were acquired on a Nikon Eclipse Ti-2 inverted microscope.

### Satellite cell isolation by fluorescence activated cell sorting

Cryobanked tongue tissue was thawed and enzymatically dissociated using collagenase IA, Dispase II, and DNase with mechanical disruption using a gentleMACS dissociator and serial trituration. After filtering, red blood cells were lysed and cells were resuspended in FACS buffer (1% BSA, 2.5% FBS, 2 mM EDTA, all in PBS). Cells were stained with antibodies against CD324, CD45, CD82, and CD56, together with live-dead fixable blue discrimination dye or DAPI. Sorting was performed on a Cytek Aurora or BD Aria Fusion using a 100 μm nozzle. Satellite cells were defined as live CD324-/CD45-/CD82+/CD56+ cells.

### Satellite cell proliferation assays

For proliferation assays, 1,000 sorted satellite cells were plated on collagen-coated 8-well chamber slides in proliferation medium consisting of DMEM/F12, 20% fetal bovine serum, GlutaMAX, and antibiotic-antimycotic. On the day after plating, medium was refreshed and supplemented with 2.5 ng/mL FGF2. On Day 4, cells were pulsed with 10 μM EdU for 1 h, fixed, permeabilized, and processed using the Click-iT Plus EdU imaging kit according to the manufacturer’s instructions. Nuclei were counterstained with DAPI. Images were acquired on a Nikon Ti-2 inverted microscope.

### Differentiation and fusion assays

For differentiation assays, 1,000 sorted satellite cells were plated on collagen-coated 96-well plates in proliferation medium. Medium was replaced every other day, with 2.5 ng/mL FGF2 added on Days 2, 4, and 6. On Day 7, cells were switched to differentiation medium containing DMEM/F12, 5% horse serum, GlutaMAX, and antibiotic-antimycotic and cultured without further medium changes for 3 days. On Day 10, cells were fixed and stained for MYOGENIN or MyHC using the MF20 antibody. Fusion index was calculated as the number of nuclei within MyHC-positive myotubes containing at least two nuclei divided by the total number of nuclei. MyHC-positive area was quantified in FIJI by threshold-based measurement after image scaling.

### Bulk RNA Sequencing

For transcriptomic analysis, 1,000 satellite cells from each control, IC2 LOM, and pUPD11 sample were expanded for 7 days and differentiated for 2 days prior to harvest on Day 9. RNA was isolated using the Maxwell RSC simplyRNA Tissue Kit. RNA quantity and quality were assessed by NanoDrop and Agilent Bioanalyzer. Five nanograms of total RNA was used for library preparation with the NextFlex Rapid Directional RNA-seq kit v2.0 and the RiboNaut rRNA depletion kit. Paired-end libraries were sequenced on a NovaSeq X platform with a target depth of 50 million reads per sample.

### RNA-seq analysis

FASTQ files were quality checked ^33^, aligned to the hg38 reference genome using HISAT2 ^29^, and quantified with featureCounts ^30^. Differential expression analysis was performed using DESeq2 ^31^. Differentially expressed genes were analyzed using a hypergeometric test and the Benjamini & Hochberg method. Genes of interest were displayed as either log2 fold change relative to control or filtered based on specific gene lists and displayed as volcano plots ^34^. The volcano plots were used to agnostically display the significant differentially expressed genes enriched in the selected pathway. The genes for the selected pathway were obtained from the Molecular Signature Database for Gene Set Enrichment Analysis (GSEA) by Broad Institute. The GSEA human gene lists M3744 (GOBP_NOTCH_SIGNALING_PATHWAY), M15486 (GOBP_SKELETAL_MUSCLE_CELL_DIFFERENTIATION), and M5909 (HALLMARK_MYOGENESIS) were used. The volcano plots display genes with p value < 0.01 and log2 fold change >0.5. Batch correction was completed on comparisons involving IC2 LOM samples.

### Pathway and network analysis

Differentially expressed genes meeting thresholds of p <0.05 and absolute log2 fold change ≥ 0.5 were analyzed using Enrichr with the Reactome Pathways 2024 library. Terms were ranked by combined score ^32^. STRING database analysis was performed using the same input gene sets, with all evidence sources included, high-confidence interaction score, and no more than 10 first-shell interactors.

### Proliferation during differentiation

For differentiation time-course assays, satellite cells were plated in triplicate and switched to differentiation medium as described above. At defined intervals corresponding to days 8, 9, and 10 of the overall culture timeline, cultures were pulsed with 10 μM EdU for 2 h, fixed, and processed for EdU incorporation. Cells were then stained for MyHC. Proliferation was quantified as the fraction of EdU-positive nuclei among total nuclei, and in a secondary analysis as the fraction of EdU-positive nuclei within or associated with MyHC-positive fused regions.

### Quantification and statistical analysis

All image analyses were performed in FIJI unless otherwise noted. Statistical analyses were performed in GraphPad Prism v10.6.1. Comparisons among three groups were analyzed by one-way ANOVA, and distribution analyses were performed by two-way ANOVA as indicated in the figure legends. Exact n values, the meaning of each data point, and statistical tests are specified in the corresponding figures and legends. P values are shown in the figures.

## Acknowledgments

We thank the patients and families who contributed samples for this study. We thank Meagan M.A. Wu for the tongue schematic used in Figure 1A. We thank Shannon V. Tringola for blinded analysis support, the CHOP Research Institute Center for Applied Genomics for assistance with very low-input RNA sequencing, Dr. Prabhat K. Sharma and Dr. Stephan A. Grupp for assistance with spectral flow cytometry, and the Penn Center for Musculoskeletal Disorders histology core. The graphical abstract was created in BioRender (agreement number YF29LLTO9W; https://BioRender.com/yof1m3x).

This work was supported by the PCMD pilot grant P30 AR069619, CHOP Academic Enrichment Funding, the Cornerstone Beckwith-Wiedemann syndrome research fund, the Victoria Fertitta Fund as part of the Lorenzo Turtle Sartini Jr. Endowed Chair in Beckwith-Wiedemann Syndrome Research, and NIH R01DE033646 to J.M.K.

## Author contributions

Conceptualization, E.D.T. and J.M.K.; Methodology, E.D.T., A.T.N., M.A.B., and J.M.K.; Investigation, E.D.T., A.T.N., and M.A.B.; Formal Analysis, E.D.T., A.T.N., M.A.B., R.D.P., G.K.-S., M.F., and S.N.; Resources, D.K. and H.P.K.; Writing – Original Draft, E.D.T. and J.M.K.; Writing – Review & Editing, all authors; Supervision, J.M.K.; Funding Acquisition, J.M.K.

## Declaration of generative AI and AI-assisted technologies in the writing process

During the preparation of this work the authors used CHOP gpt5.2 in order to improve manuscript flow. After using this tool, the authors reviewed and edited the content as needed and take full responsibility for the content of the published article.

## Summary Figure

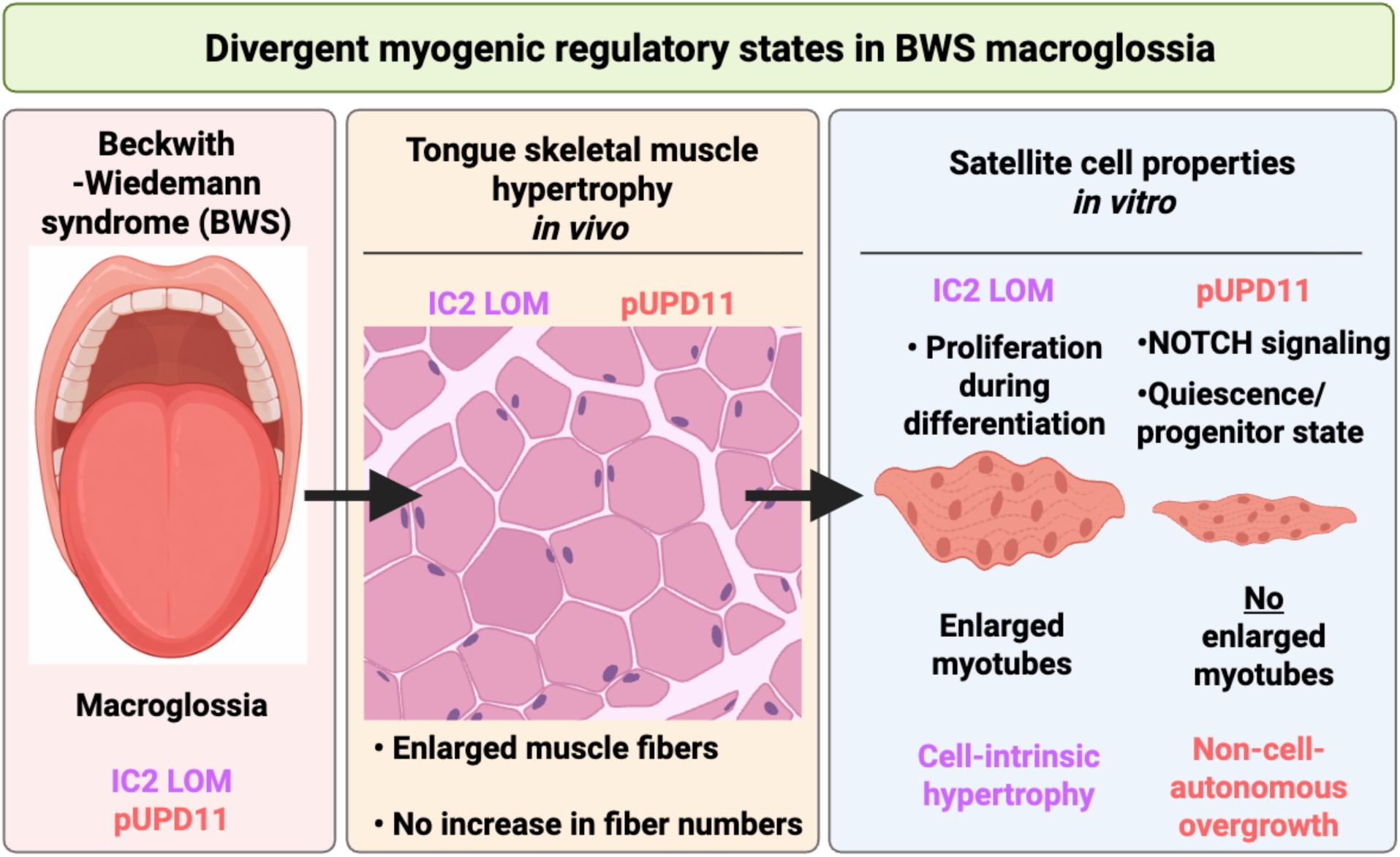

