## Supplementary for "Myogenic dysregulation underlies tongue overgrowth in Beckwith-Wiedemann syndrome"

This PDF file includes:

Figures S1–S8

### Supplemental Figures

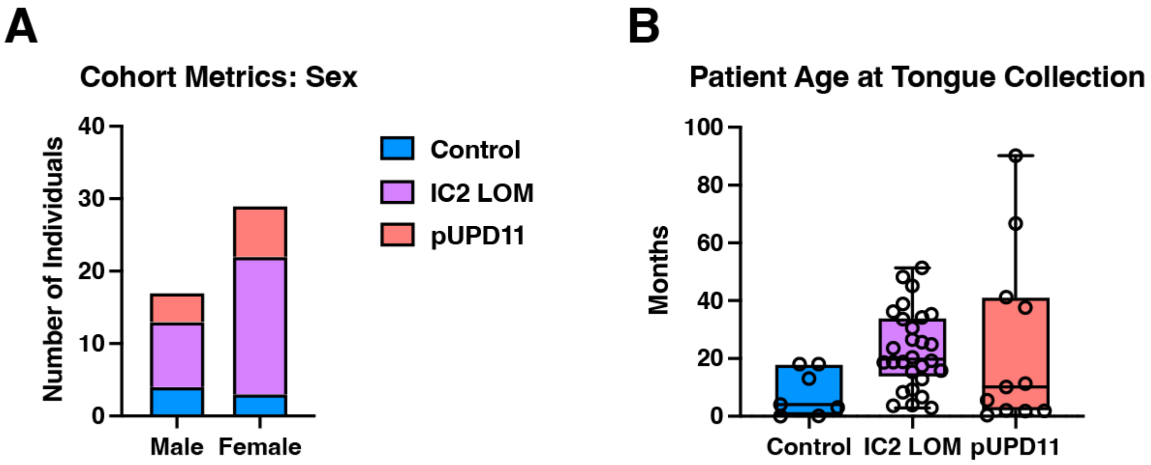


**Figure S1. Patient cohort characteristics for histological analyses.**

(A) Sex distribution of tongue specimens from control/non-BWS (blue), IC2 LOM (purple), and pUPD11 (salmon) cohorts.

(B) Age of patients at the time of tongue specimen collection. Each data point represents a specimen from a unique patient.

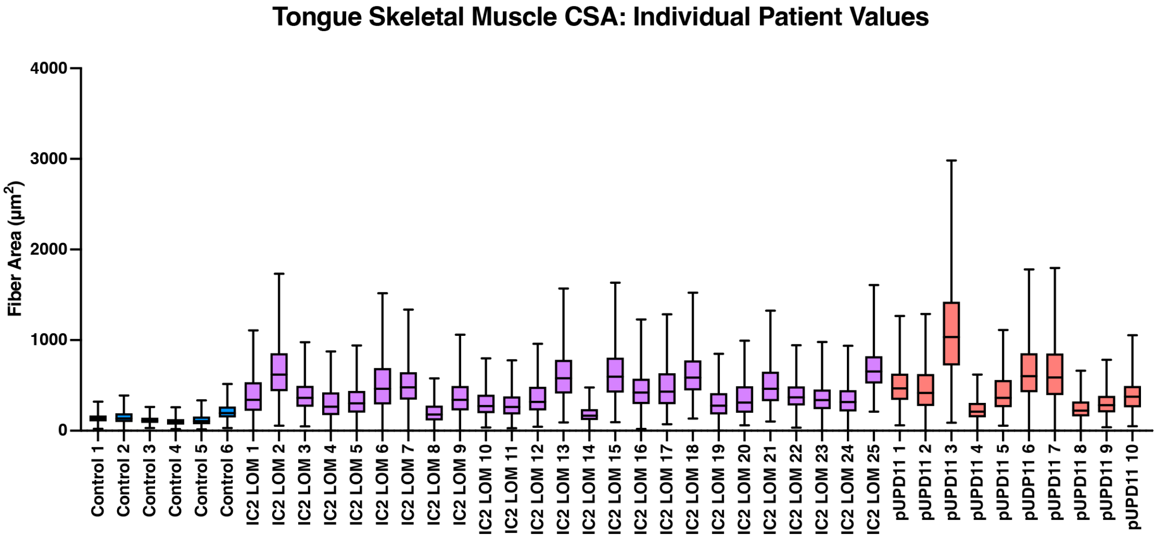


**Figure S2. Skeletal muscle fiber cross-sectional area by individual patient.**

Patient-level breakdown of muscle fiber cross-sectional area (CSA) measurements corresponding to aggregated data shown in Fig. 1C.

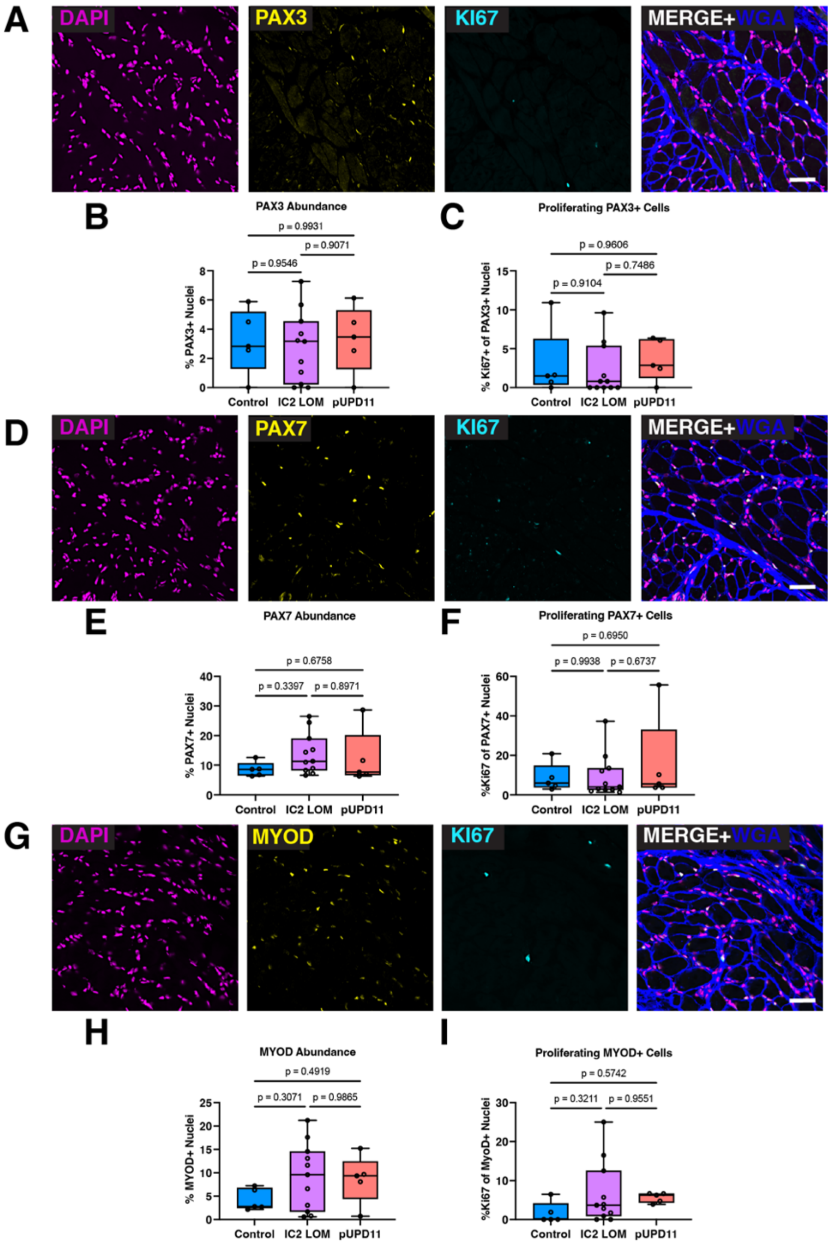
A
DAPI
PAX3
KI67
MERGE+WGA
B
PAX3 Abundance
p = 0.9931
p = 0.9546
p = 0.9071
% PAX3+ Nuclei
Control
IC2 LOM
pUPD11
C
Proliferating PAX3+ Cells
p = 0.9606
p = 0.9104
p = 0.7486
% KI67+ of PAX3+ Nuclei
D
PAX7
E
PAX7 Abundance
p = 0.6758
p = 0.3397
p = 0.8971
% PAX7+ Nuclei
F
Proliferating PAX7+ Cells
p = 0.6950
p = 0.9938
p = 0.6737
%KI67 of PAX7+ Nuclei
G
MYOD
H
MYOD Abundance
p = 0.4919
p = 0.3071
p = 0.9865
% MYOD+ Nuclei
I
Proliferating MYOD+ Cells
p = 0.5742
p = 0.3211
p = 0.9551
%KI67 of MyoD+ Nuclei

**Figure S3. Comparable abundance and proliferation of myogenic precursors in control and BWS tongues.**

**(**A) Representative cryosection images showing nuclei (DAPI; magenta), PAX3 (yellow), KI67 (cyan), and wheat germ agglutinin (WGA; blue) marking myofiber boundaries. Scale bar: 50 μm.

(B) Quantification of $PAX3^+$ nuclei in control (n = 4), IC2 LOM (n = 11), and pUPD11 (n = 5) tongue samples.

(C) Quantification of $KI67^+$ cells among $PAX3^+$ nuclei.

(D) Representative images showing nuclei (DAPI; magenta), PAX7 (yellow), KI67 (cyan), and WGA (blue). Scale bar: 50 μm.

(E) Quantification of $PAX7^+$ nuclei.

(F) Quantification of $KI67^+$ cells among $PAX7^+$ nuclei.

(G) Representative images showing nuclei (DAPI; magenta), MYOD (yellow), KI67 (cyan), and WGA (blue). Scale bar: 50 μm.

(H) Quantification of $MYOD^+$ nuclei.

(I) Quantification of $KI67^+$ cells among $MYOD^+$ nuclei.

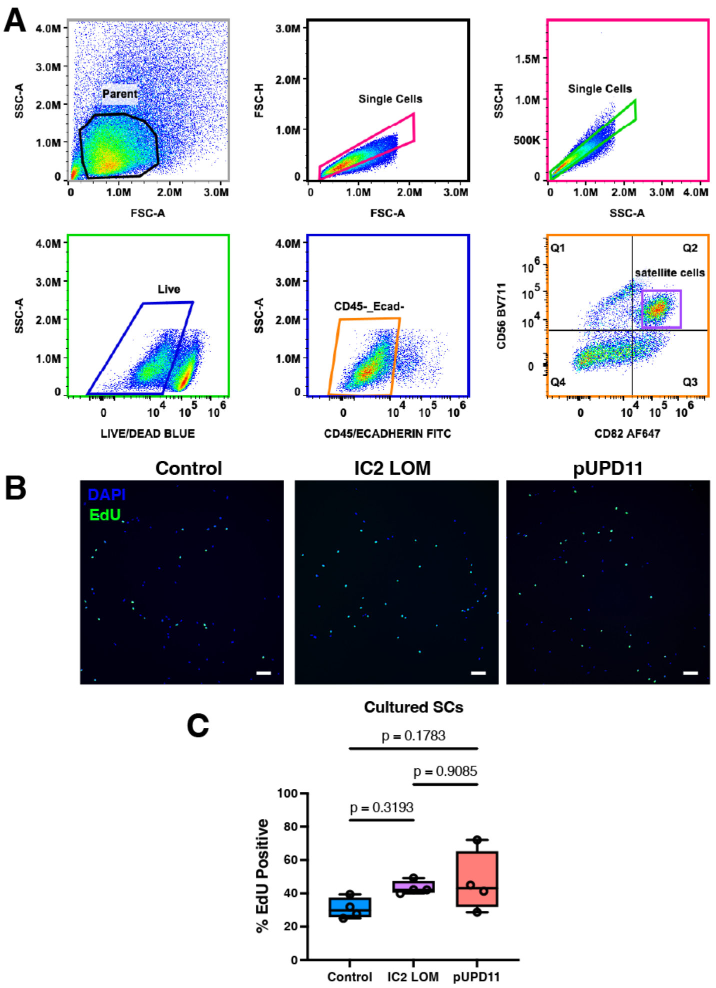


**Figure S4. Prospective isolation and *in vitro* proliferation analysis of human satellite cells.**

(A) FACS gating strategy for satellite cell isolation. Live satellite cells were gated and defined as CD324-/CD45-/CD82+/CD56+ (ECAD-/LCA-/TSPAN27+/NCAM+).

(B) Representative images of control, IC2 LOM, and pUPD11 satellite cells stained for EdU incorporation (green) on day 4 post-plating following a 1-h EdU pulse. Scale bar: 100 µm.

(C) Quantification of $EdU^{+}$ nuclei normalized to total nuclei. Each data point represents cells derived from one unique patient. Statistical significance was determined by ANOVA.

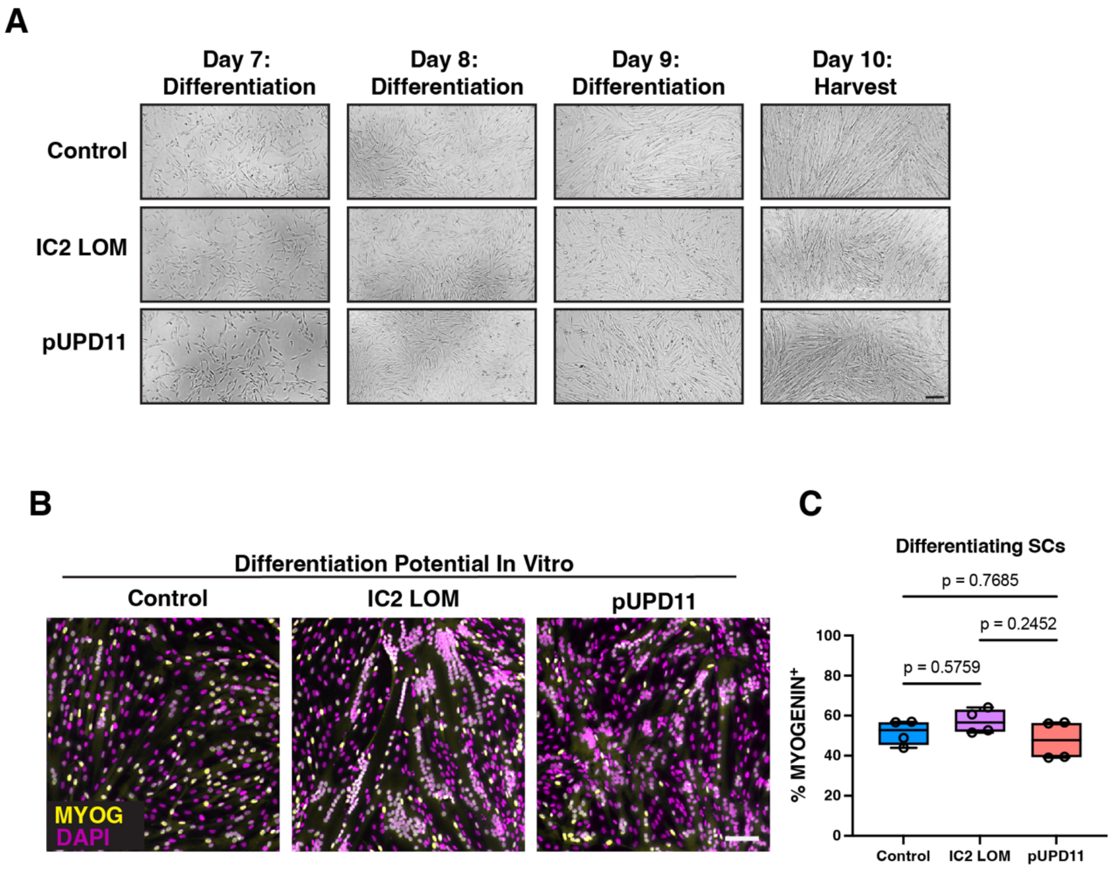


**Figure S5. Satellite cell differentiation properties in vitro.**

(A) Phase contrast images of satellite cells grown at the indicated time points. Differentiation was induced on day 7 post-plating by switching to differentiation medium. Scale bar: 100μm.

(B) Representative images of satellite cells fixed at day 10 post-plating (3 days of differentiation) and stained with MYOGENIN (MYOG; yellow) and nuclei (DAPI, magenta).

(C) Quantification of MYOGENIN$^+$ nuclei. Each data point represents one unique patient (n = 4 per cohort). Significance was determined by ANOVA.

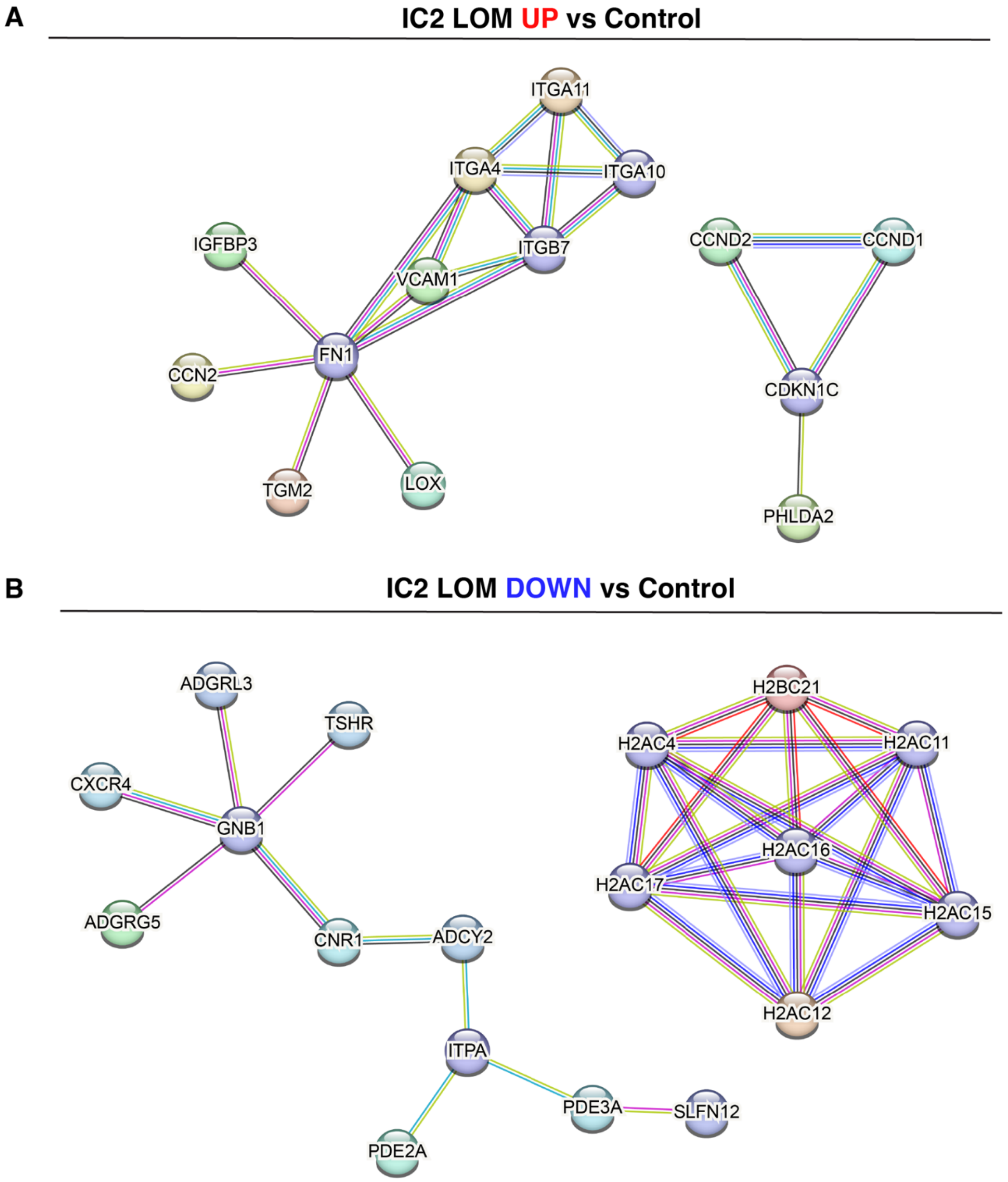


**Figure S6. STRING network analysis of protein nodes implied from differentially expressed genes in IC2 LOM versus control satellite cells.**

(A) Interaction network of genes upregulated in IC2 LOM samples at day 9 post-plating (day 2 of differentiation).

(B) Interaction network of genes upregulated in control samples relative to IC2 LOM. Only nodes with ≥4 interactors are shown.

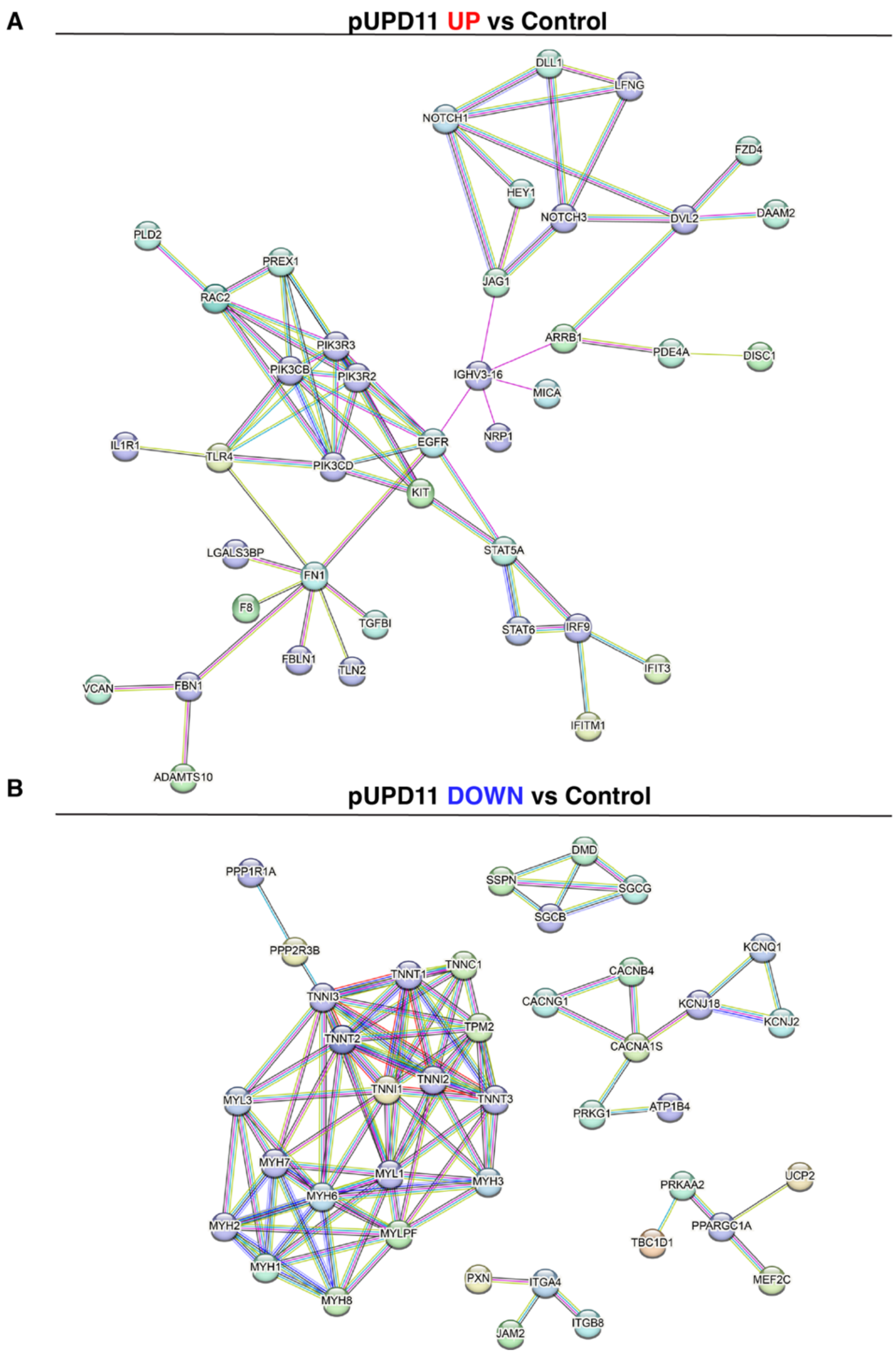


**Figure S7. STRING network analysis of protein nodes implied from differentially expressed genes in pUPD11 versus control satellite cells.**

(A) Interaction network of genes upregulated in pUPD11 samples at day 9 post-plating (day 2 of differentiation).

(B) Interaction network of genes upregulated in control samples relative to pUPD11. Only nodes with ≥4 interactors are shown.

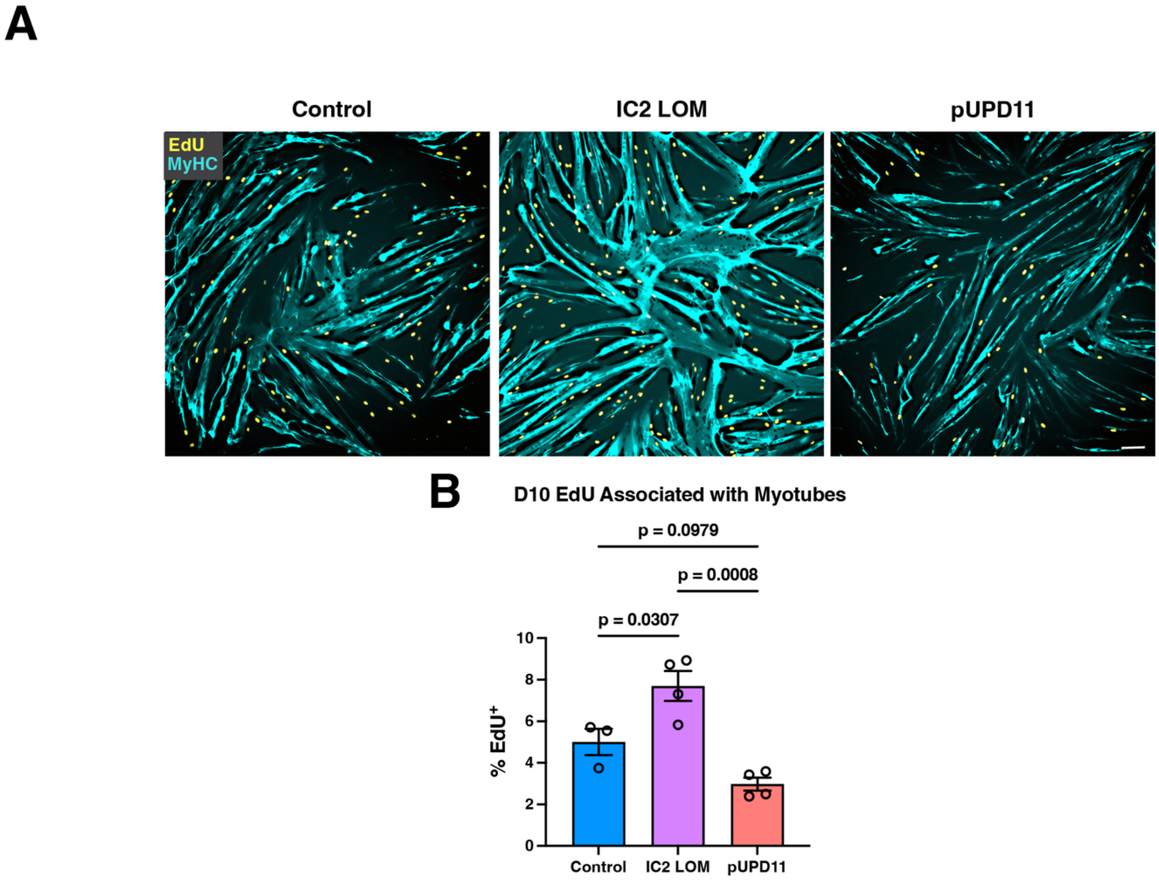


**Figure S8. Proliferative nuclei associated with myotubes during differentiation.**

(A) Representative images of control, IC2 LOM, and pUPD11 satellite cells pulsed with EdU for 2 h prior to fixation on day 10 (day 3 of differentiation). Scale bar: 100 µm. Images correspond to those shown in Fig. 5F with the DAPI channel removed for clarity.

(B) Quantification of $EdU^+$ nuclei located within or immediately adjacent to $MYHC^+$ myotubes. Each data point represents cells derived from one unique patient. Statistical significance was determined by ANOVA.
